# Deciphering the Sequence Basis and Application of Transcriptional Initiation Regulation in Plant Genomes Through Deep Learning

**DOI:** 10.1101/2025.04.22.649999

**Authors:** Pengfei Gao, Lijie Lian, Wanjie Feng, Yuxue Ma, Jieni Lin, Liya Qin, Shanmeng Hao, Haonan Zhao, Xuantong Liu, Jing Yuan, Zongcheng Lin, Xia Li, Yuefeng Guan, Xutong Wang

## Abstract

Transcription initiation is a critical regulatory step in plant gene expression, yet its sequence determinants remain largely elusive. Here we introduce *GenoRetriever*, an interpretable deep learning model that deciphers the sequence basis of transcriptional initiation regulation across plant genomes. Trained on STRIPE-seq data from 16 soybean tissues and six other crop species, *GenoRetriever* identifies 27 core sequence motifs that govern transcription start site (TSS) selection and usage. The model predicts TSS locations and usage levels with high accuracy, as validated by in silico motif insertions, saturation mutagenesis, and CRISPR-Cas9 promoter editing. It further reveals that 31.85% of natural variation between wild and domesticated soybean drives shifts in promoter motif usage during domestication, and uncovers lineage-specific motif effects between monocots and dicots. This interpretable model and its user-friendly web server for promoter analysis and design make GenoRetriever both a methodological innovation and practical tool for plant functional genomics and crop improvement.

## Introduction

Transcriptional initiation is a critical regulatory layer in gene expression, fundamentally determining transcript abundance and influencing cellular and organismal functions^1–4^. Precisely regulating gene expression is particularly significant for improving crop traits, as precisely regulating gene expression could enhance plant plasticity in yield and their adaptation, as controlled gene expression adjustments can yield desirable agronomic outcomes^5^. Promoter editing, for instance, enables targeted regulation of gene expression levels rather than complete gene knockouts, avoiding extreme phenotypic changes. An example in soybean shows that precise modulation of *GmNARK* expression to moderately increase nodulation can enhance yield, unlike knockout mutants exhibiting excessive nodulation without yield benefits^6^. Similarly, in rice, targeted regulatory sequence mutations in *IPA1* significantly increased grain yield by decoupling panicle number and size^5^.

High-resolution experimental methods like Cap Analysis Gene Expression (CAGE), nanoCAGE, and Survey of TRanscription Initiation at Promoter Elements with high-throughput sequencing (STRIPE-seq) have significantly improved our ability to map transcription start sites (TSS) at single-base resolution^7–10^. Concurrently, artificial intelligence (AI)-based models, including Basenji and Enformer^11, 12^, have substantially enhanced the prediction of gene expression levels. Despite these advances, current AI methods primarily function as “black boxes,” limiting our understanding of the specific nucleotide-level determinants governing transcription initiation. A notable gap remains in identifying and understanding additional sequence motifs beyond classical elements (e.g., TATA-box, initiator elements) crucial for transcription initiation in plants. Several important questions remain unanswered, including why bidirectional transcription initiation is prevalent in vertebrates yet uncommon in plants, and what biological consequences this difference entails. Additionally, it remains unclear how evolutionary and domestication processes have shaped the conservation or divergence of these regulatory sequences between monocots and dicots.

To address these critical questions, we conducted a systematic multi-species study. We expanded a validated, modified STRIPE-seq protocol originally used in soybean to generate high-resolution TSS profiles across 12 tissues from cultivated soybean (*Glycine max*, Williams 82), wild soybean (*Glycine soja*, PI468916), and the leaves tissues from selected dicots including cotton (*Gossypium spp*.), rapeseed (*Brassica napus*), tomato (*Solanum lycopersicum*), and monocots such as maize (*Zea mays*), wheat (*Triticum aestivum*), and rice (*Oryza sativa*). Complementing this experimental approach, we developed a novel interpretable deep-learning framework, *GenoRetriever*, designed to accurately predict TSS locations and transcriptional abundance while elucidating sequence contributions to transcription initiation. Our integrative analysis provides novel insights into plant transcription initiation mechanisms and identifies new cis-elements essential for gene regulation. This study represents a significant advancement in our understanding of plant gene expression regulation and provides valuable tools for crop improvement and functional genomic applications.

## Results

### Base-pair resolution interpretation of sequence determinants and precise prediction of transcription initiation in soybean

To ensure highly accurate transcription start site (TSS) annotations, we reanalyzed previously validated STRIPE-seq data from eight soybean tissues, including five vegetative (leaves, shoot, stem tips, roots, and mature nodules) and three reproductive (flowers, pods, and seeds)^10^ (**Fig. 1a, Supplementary Table 1)**. Approximately 40,000 reliable TSS regions (TSRs) per tissue were annotated using the telomere-to-telomere (T2T) reference genome of cultivated soybean (Williams 82)^13^. From these data, we extracted 4,650 bp sequences containing dominant TSSs located within key regulatory regions (proximal promoters and 5′ UTRs). Each sequence spanned 4,000 bp centered on the TSS (3,500 bp upstream and 500 bp downstream), with an additional 325 bp buffer at both ends to capture flanking context while reducing edge noise (**Fig. 1a)**. We then developed *GenoRetriever*, a deep learning framework comprising three modules: a motif consensus network, a supplementary effect consensus network, and a feature prediction network (**Fig. 1b, Extended Data Fig. 1**). The motif consensus network, inspired by Puffin^14^, employs a two-layer convolutional architecture for sequence feature extraction and encoding **(Extended Data Fig. 1**). To identify robust sequence features, we applied artificial knowledge distillation by training 15 independent networks on STRIPE-seq data from each tissue and subsequently analyzing the correlations among convolutional kernels **(Extended Data Fig. 1**). A connectivity graph based on correlation coefficients was used to pinpoint biologically meaningful consensus patterns, which were categorized according to their encoding-layer response curves (**Fig. 1b)**. These patterns fell into two major groups: canonical motifs and initiator elements (Inr). Inr elements exhibited sharply localized effects, while the motifs showed broader influence ranges (**Fig. 1c, Extended Data Fig. 2**). Furthermore, Inr patterns were subdivided into short and long Inr based on their effect spread. Alignment against the JASPAR database ^15^confirmed numerous known elements; this approach yielded five short Inr sequences, four long Inr sequences, and 18 motifs, including nine known (e.g., TATA-box, YY1, DREB1) and nine unknown motifs, thereby significantly advancing our understanding of transcription initiation in plants (**Fig. 1c, Extended Data Fig. 2**).

**Figure 1.**
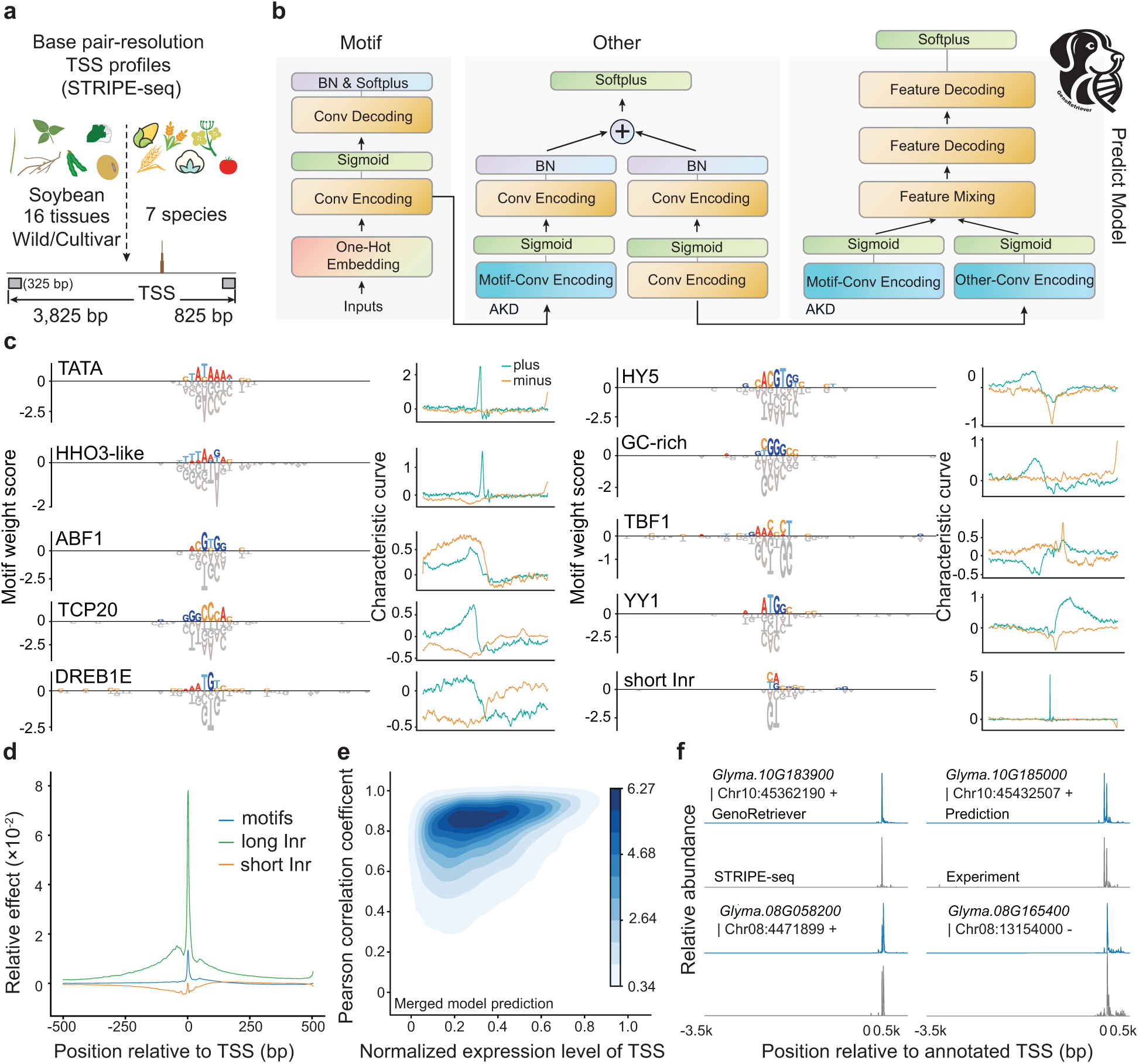
Overview of the *GenoRetriever* model and its performance in interpreting and predicting transcriptional initiation. (a) Training data comprised base-pair resolution TSS profiles from STRIPE-seq in 16 tissues of wild and cultivated soybean and in leaf tissue of multiple crops including cotton, rice, maize, wheat, tomato and rapeseed. For each TSS, a 4,325 bp window spanning 3,825 bp upstream to 500 bp downstream was used. (b) Workflow for *GenoRetriever* training and prediction. Two consensus networks first extract important sequence patterns by artificial knowledge distillation to generate weight matrices for each pattern. These matrices are then loaded into the prediction network to accurately forecast transcription initiation signals. (c) Ten representative sequence patterns out of the 27 identified by the motif consensus network, together with their characteristic response curves plotted over a 1-kb window centered on each pattern. (d) Average effect profiles of motifs and initiator elements at single TSSs. The x-axis shows relative position around the TSS (0 marks the TSS) and the y-axis shows the log scaled effect magnitude. (e) Density estimates of prediction accuracy versus relative expression level for the multi-tissue Williams 82 model. Accuracy is measured by the Pearson correlation coefficient. The x-axis shows normalized TSS expression and the y-axis shows correlation values. (f) Example comparison of predicted and observed transcription initiation signal profiles on the test set for the multi-tissue Williams 82 model.

The feature prediction network was constructed using a U-Net-like sequence-to-sequence model enhanced with ResNet-style residual connections and a custom Multi-head Efficient Channel Attention (MECA) mechanism **(Extended Data Fig. 1**). Our *GenoRetriever* model, which implements this architecture, ultimately identified 27 distinct sequence patterns. Most motifs exhibited unidirectional effects favoring the plus strand, with the exception of DREB1E, which showed bidirectional influence, potentially explaining the scarcity of bidirectional promoters in soybean^10^ (**Fig. 1c, Extended Data Fig. 2**). Comparative analyses, including ablation experiments, indicated that transcription initiation is jointly determined by motifs and Inr elements, with both contributing comparably to TSS expression levels. On average, individual motifs had a limited influence, significantly weaker than that of Inr sequences. Within the Inr components, long Inr elements exerted stronger effects than short Inr (**Fig. 1d)**. While motifs functioned through diverse and broadly distributed interactions near TSS regions, Inr elements displayed more fixed positions and quantities at individual loci (**Fig. 1d)**. These findings suggest that transcription initiation in soybean is governed by combinatorial interactions between Inr sequences and one or more motifs (**Fig. 1d)**.

After extracting the supplementary effects, the 27 identified sequence patterns were integrated with an additional supplementary effect layer into the feature prediction network as fixed weights (**Fig. 1b)**. The model was trained on the merged STRIPE-seq dataset from all eight tissues of Williams 82 to predict TSS expression levels. Based on the 27 sequence patterns and supplementary features, our prediction model achieved an average Pearson correlation of 75.25% on the test set (**Fig. 1e)**. Notably, the model’s predictive accuracy was positively associated with the relative expression abundance of TSSs: TSSs with predictive correlations below 0.40 generally had low expression levels (relative abundance ≤ 0.40), whereas TSSs with relative abundance above 0.40 exhibited predictive correlations mostly around 0.80 (**Fig. 1e)**. These results indicate that our model reliably predicts transcription initiation activity in soybean and provides a solid foundation for further functional dissection of sequence features. For example, predictions the TSS for representative genes (*Glyma.10G183900* and *Glyma.10G185000*) closely matched the STRIPE-seq profiles, demonstrating the accuracy and robustness of our approach (**Fig. 1f)**.

### Accurate interpretation of motif effects on transcription initiation abundance and localization

Motifs are key cis-regulatory elements in gene promoters that can be activated by transcription factor (TF) binding or repressed when such binding is absent (**Fig. 2a)**. Because the first convolutional layer in our prediction model derives its weights from the consensus network’s sequence features, we can “virtually switch off” individual motif effects by suppressing the corresponding convolutional kernel activations **(Extended Data Fig. 1**). This *in silico* motif ablation enables us to evaluate the impact of each motif on TSS signal intensity and position. In this analysis, we defined a 1-kb window (500 bp upstream and downstream of the TSS) and measured the effect of motif knockout on TSS signal abundance. By subtracting the predicted signal values before and after motif ablation at single-base resolution along the window, we generated an effect curve that quantifies the influence of each motif on TSS signal (**Fig. 2b)**. Using this approach, we observed distinct modes of action among different motifs. For example, TCP20 promoted transcription surrounding TSS, whereas DREB1E functioned predominantly as repressors. Additionally, some motifs, exemplified by the TATA-box, displayed a dual effect by repressing signals immediately adjacent to the TSS while enhancing transcription exactly at the TSS (**Fig. 2b)**. Notably, several newly identified motifs also demonstrated significant regulatory effects, underscoring the importance of uncovering these elements to fully decipher TSS regulation (**Fig. 2b)**.

**Figure 2.**
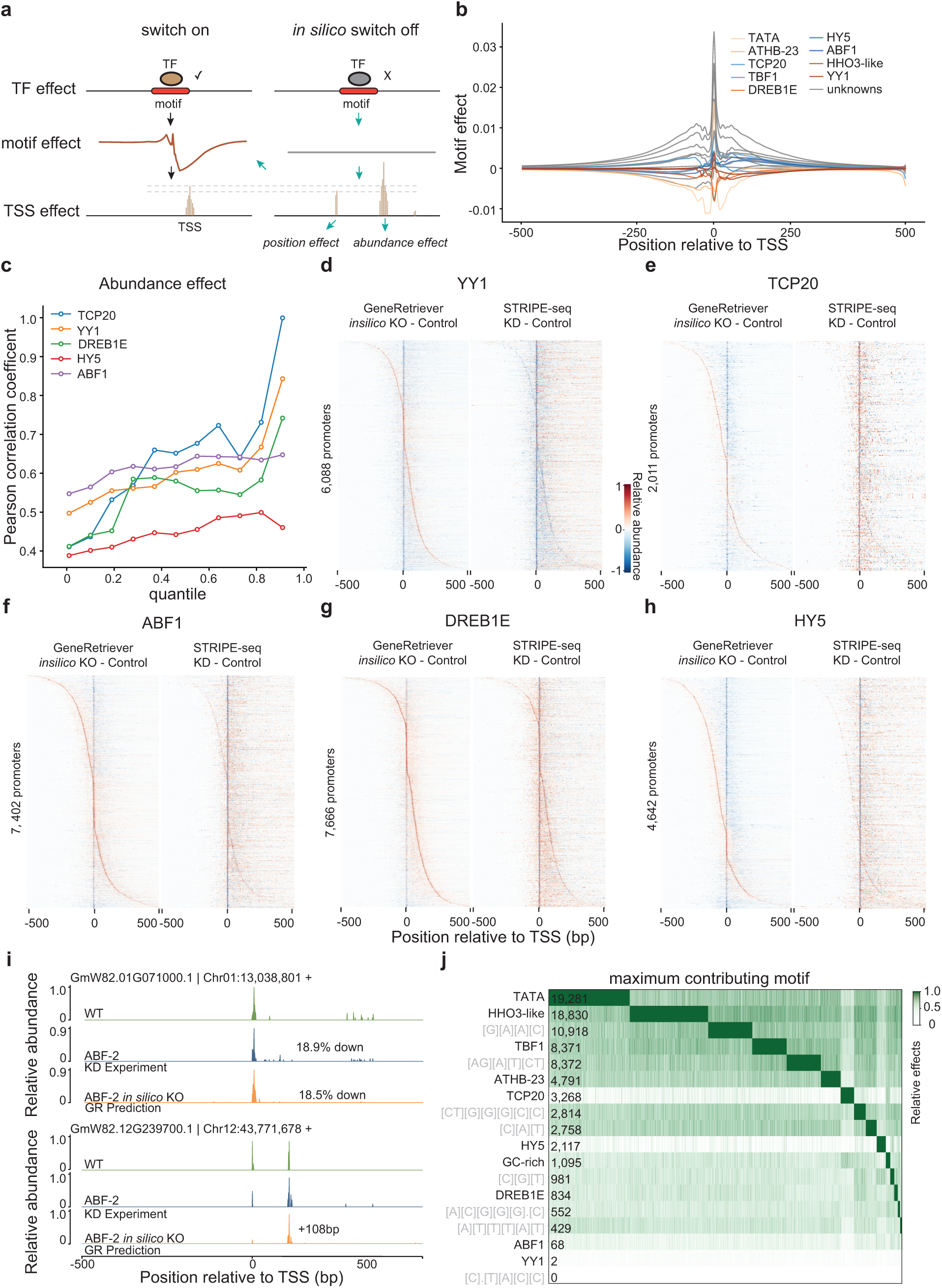
Dissecting motif effects on transcription initiation in soybean using *GenoRetriever*. **a** Workflow for *in silico* motif ablation to quantify motif effects on TSS positioning and abundance. *GenoRetriever* switches off the weight for a given motif and measures the resulting change in the predicted initiation signal. **b** Average effect curves of each motif on TSS signal. The y-axis shows effect magnitude and the x axis shows position relative to the TSS (0 marks the TSS). **c** Correlation between in silico knockout predictions and *in vivo* transcription factor knockdown experiments for five motifs. The y-axis shows Pearson correlation coefficients and the x-axis shows quantiles of TSSs ranked by predicted motif effect on abundance. **d–h** Heatmaps of positional effects for five motifs: YY1 (d), TCP20 (e), ABF1 (f), DREB1E (g) and HY5 (h). Each row is one promoter, the x-axis is position relative to the TSS and cell color represents the change in signal abundance after *in silico* motif knockout (KO) or *in vivo* factor knockdown (KD). **i** Example of abundance and positional effects for the ABF1-2 motif on gene *GmW82.01G071000*. The plot compares in vivo ABF1-2 knockdown (KD) with *in silico* motif knockout (KO) predictions over a 1-kb window centered on the TSS. **j** Promoter classification by dominant motif effect. Columns represent promoters and rows represent motif types. Numbers indicate how many promoters rely on each motif as the primary contributor to transcription initiation.

To further evaluate the influence of motifs on transcriptional activity, we selected four motifs (DREB1E, YY1, ABF1, and HY5) due to the well-characterized nature of their binding transcription factors. For each motif, we quantified the relative change in TSS signal intensity within a ±200 bp window before and after *in silico* motif ablation. In parallel, we performed experimental validations via knockdown experiments targeting TFs associated with five motifs (YY1, TCP20, ABF1, DREB1E, and HY5) **(Supplementary Table 1)**. These *in vivo* knockdown studies, conducted using a hairy-root transformation system to avoid the transcriptional shock common to knockout approaches in soybean, enabled us to compare changes in TSS abundance between transgenic (RNAi-mediated knockdown) and wild-type samples **(Extended Data Fig. 3**). Correlating the *in silico* knockout predictions with the experimental data, we observed a strong positive relationship (**Fig. 2c)**; the predictive accuracy (measured as correlation coefficients) increased as the modeled motif effect on TSS abundance became more pronounced (**Fig. 2c)**. This finding demonstrates that our *GenoRetriever* model not only accurately forecasts the quantitative impact of motif loss on TSS signal but also reinforces the abundance regulatory importance of these motifs. Interestingly, our analysis revealed that motif effects are not limited to TSS abundance, they also modulate the precise positioning of TSSs. For example, based on the effect curve for the TATA-box, we noted that its regulatory impact extends to positioning control. To evaluate this hypothesis, we selected promoters containing motifs within a window from 2,000 bp upstream to 500 bp downstream of the annotated TSS. Promoters were sorted according to the positional shift in the maximum predicted signal following motif ablation, and these shifts were compared against experimental STRIPE-seq data (**Fig. 2d-h)**. In the experimental group, convolution-based smoothing using a window size of [5, 3] accentuated these shifts. Both the *in silico* predictions and the experimental results revealed that the deletion of a motif can lead to a discernible shift in the position of the maximum TSS signal, suggesting that these motifs contribute to both the magnitude and localization of transcription initiation (**Fig. 2i)**.

Finally, to comprehensively quantify the overall effect of each motif on TSS regulation, we employed a Fourier transform-based squared difference approach that simultaneously captures changes in TSS abundance and positional effects. Using the resulting effect scores, we classified promoters in Williams 82 according to the dominant motif effect (**Fig. 2j, Supplementary Table 2)**. Intriguingly, while many promoters are primarily regulated by the TATA-box, a substantial subset is predominantly influenced by other motifs. For example, TATA-box regulation was observed in a large fraction (25.01%) of promoters, whereas promoters dominated solely by the YY1 motif were rare. This distribution may reflect cooperative interactions among motifs, as suggested by the YY1 element (**Fig. 2j)**. In summary, our integrated *in silico* and experimental analyses demonstrate that motifs exert dual regulatory roles on both the abundance and positioning of TSSs in soybean. These findings not only validate the predictive power of the *GenoRetriever* model but also deepen our understanding of the cis-regulatory mechanisms that shape transcription initiation at base-pair resolution.

### Effect shift of motifs caused by natural variation during domestication

Cultivated soybean was domesticated from wild soybean approximately 5,000 to 9,000 years ago^16,17^. To investigate how key motif regulatory patterns have changed during the evolution of soybean, we assembled T2T genome for the wild soybean germplasm PI468916 (**Fig. 3a)**. PI468916 is an important resource for domestication studies and harbors resistance loci to soybean cyst nematode (SCN) ^18, 19^. Accurate model training for PI468916 requires a high-quality genome, and we achieved this by combining high-fidelity (HiFi) sequencing, Nanopore ultra-long reads, and Hi-C data **(Supplementary Table 3)**. The result was a complete, gapless T2T assembly that was further annotated with transcriptome sequencing data from six different tissues, leading to the identification of 62,237 genes and 52.83% transposable elements **(Supplementary Table 4)**. To decipher how sequence variation affects transcription initiation, we collected eight tissues from PI468916 at the same developmental stage as Williams 82. These samples were processed with our modified STRIPE-seq protocol to capture TSS profiles. After quality control and mapping the STRIPE-seq data to the newly assembled PI468916 genome, we identified 40,667 reliable TSRs covering 24,537 genes in PI468916 **(Supplementary Table 5)**.

**Figure 3.**
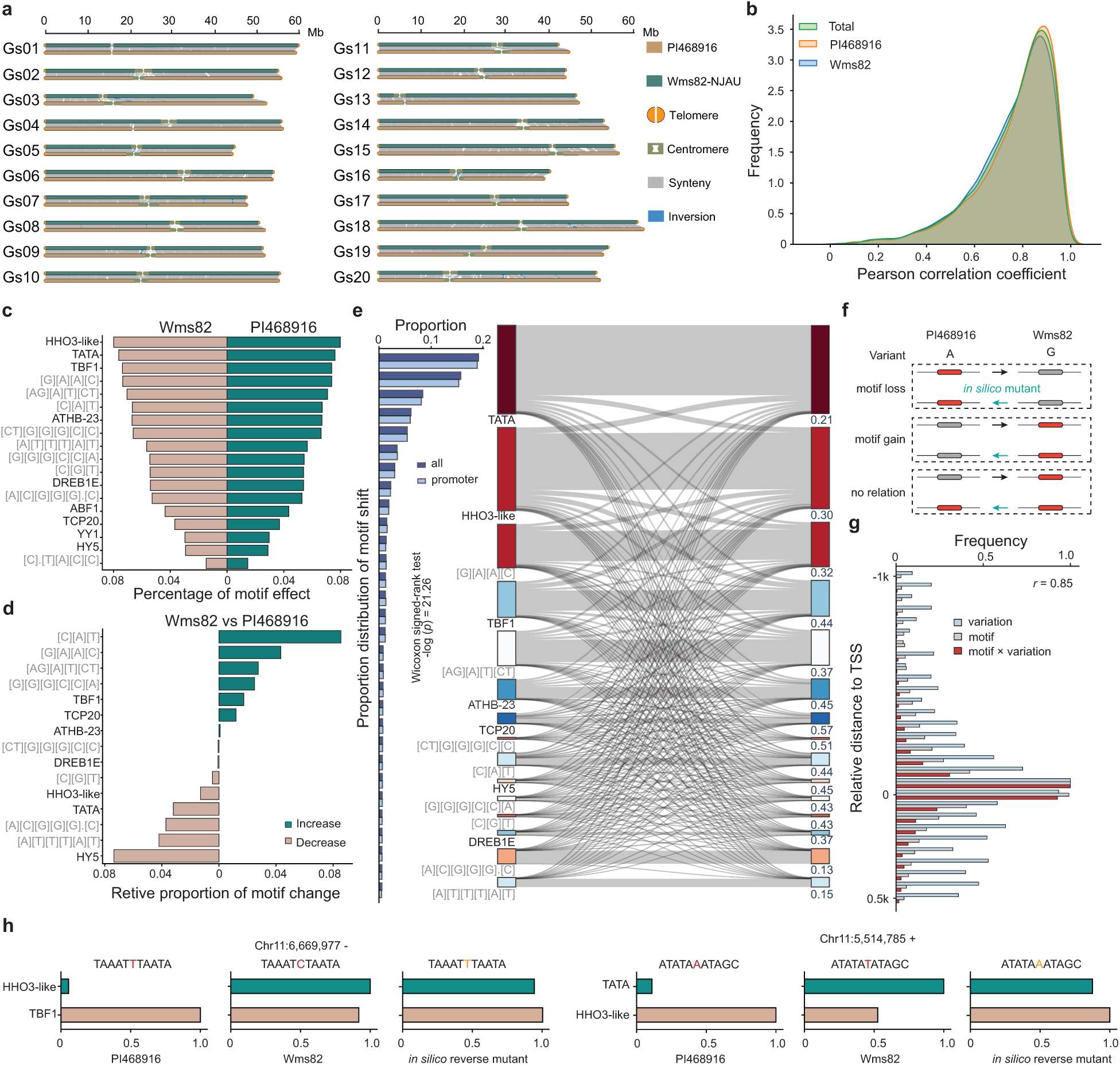
Impact of natural variation on transcription initiation during soybean domestication. **a** Telomere-to-telomere assembly of the wild soybean genome (PI468916) and synteny comparison with the cultivated Williams 82 genome. Structural variants between the two assemblies are highlighted along each chromosome. **b** Kernel density estimates of *GenoRetriever* prediction accuracy for TSS signals. Green shows the distribution of correlation coefficients for all TSS predictions pooled across both PI468916 and Williams 82. Orange shows the distribution for PI468916 alone and blue shows the distribution for Williams 82 alone. The y-axis indicates the frequency of TSSs at each correlation level and the x axis indicates Pearson correlation coefficients between predicted and observed signals. **c** Comparison of motif effect proportions between Williams 82 and PI468916. The y-axis lists motif types and the x-axis shows each motif’s share of the total regulatory effect on TSS abundance. **d** Relative changes in the number of promoters dominated by each motif between Williams 82 and PI468916. The x-axis shows the change in proportion of promoters where a given motif is dominant, and the y-axis lists the motif types. **e** Wilcoxon rank sum test comparing motif shift proportions for promoters with variants in the 2-kb upstream region versus all promoters. The y-axis lists motif shift types and the x-axis shows their proportions. The right panel shows a Sankey diagram of motif shifts from PI468916 (left) to Williams 82 (right), with labels indicating the fraction of TSSs whose dominant motif changed. **f** Illustration of the *in silico* degradation experiment used to identify critical variations that drive motif shifts. **g** Frequency distributions of motif occurrences and critical variation positions relative to the TSS. The y-axis shows distance bins from the TSS (0 at the TSS) and the x-axis shows the frequency of motif occurrences and key variations in each bin. Critical variations are those that convert the Williams 82 dominant motif to the corresponding PI468916 motif. **h** Example of a critical variation in a domestication-associated gene promoter. The y-axis lists the dominant motifs in PI468916 and Williams 82, and the x-axis shows each motif’s relative effect magnitude on transcription initiation.

Furthermore, we integrated the PI468916 STRIPE-seq data into the *GenoRetriever* model together with the Williams 82’s (**Fig. 3b & 1a)**. The model was trained on the combined datasets to learn the effects of critical variations between the two soybean types. Evaluation indicated that the model’s performance in predicting TSS signal in PI468916 was comparable to that in Williams 82, establishing a reliable foundation for motif pattern analysis between the two varieties (**Fig. 3b)**. Comparative analysis of motif effects showed that the overall weight distribution of motif effects was largely conserved between the two cultivars, with no significant changes in the absolute values (**Fig. 3c, Supplementary Table 2 & 6)**. However, when we performed a cross-species comparison of the relative proportion of each motif, we observed a symmetrical trend of subtle differences in motif effects. (**Fig. 3d)**. This result suggests that for a given promoter, the dominant motif may shift between the two soybean types. We next constructed a genome synteny relationship between PI468916 and Williams 82 and identified 39,985 pairs of TSRs for further analysis (**Fig. 3a, Supplementary Table 7)**. Promoters that aligned as the same genomic region between the two varieties were compared based on their dominant motif type. Our analysis revealed that motif shifts do indeed occur between syntenic promoters. This phenomenon was more frequently observed in promoters regulated by weaker-effect motifs, whereas promoters dominated by strong-effect motifs (such as TATA-box or HHO3-like) exhibited fewer shifts (**Fig. 3e)**.

In the PI468916 genome, we also detected 8,110,292 sequence variants specific to this wild soybean compared to the Williams 82 genome (**Fig. 3a)**. Variations located near genes provide an opportunity to better understand transcription initiation regulation during soybean domestication. To explore the link between sequence variation and motif shifts, we compared the distribution of motif shift types in promoters with sequence variants located within **2-kb** upstream and 500bp downstream of the TSS against the overall distribution of motif shifts (**Fig. 3e)**. It showed a significant difference between the two, supporting the hypothesis that sequence variation may be an important factor driving motif shift (Wilcoxon rank sum test, *p* value < 0.001). To further confirm our inference, we performed *in silico* degradation experiments. In these experiments, key variations were identified by reverting a variant nucleotide to its alternative allele. For instance, in one example, a variant where PI468916 had an “A” and Williams 82 a “G” caused the motif to shift from one type to another. When we mutated the “G” back to an “A” *in silico*, the motif shifted back accordingly (**Fig. 3f)**. Using this approach, we identified 34,226 (13.42%) variations, including 22,654 SNPs and 11,571 indels, as key drivers of the motif shift phenomenon **(Supplementary Table 8)**. We also analyzed the frequency of key variations across different positions relative to the TSS and compared this to the frequency of motif occurrence in the same regions (**Fig. 3g)**. We found a significant correlation (*r* = 0.85) was observed between the frequency of key variations and motif motif occurrence; the frequency of variations increased with distance from the TSS, peaking within 50 bp on either side (**Fig. 3g)**. This indicates that sequence variation near motifs is a major contributor to the motif shift phenomenon. In-depth analysis shows that a small number of critical sequence variations can directly lead to the disruption or creation of motif sequences within the same promoter between the two varieties (accounting for 3.70%), thus altering the relative contribution of motifs to the transcription start site (TSS) signal (**Fig. 3g)**. For example, on chromosome 11 at position 5,514,785 (negative strand), an A→T SNP disrupted the TATA-box sequence, reducing its contribution to the TSS signal and allowing an HHO3-like motif to become dominant. Conversely, at position 6,669,977 (negative strand), a T→C SNP upstream of the TSS created a new HHO3-like motif, boosting its contribution and supplanting ABF1 as the primary regulatory element (**Fig. 3h)**.

### Multi-scale sequence editing for accurate prediction of transcription initiation abundance and its applications

Although we have already demonstrated *GenoRetriever*’s excellent performance in predicting STRIPE-seq signals, it remains essential to assess whether the model can accurately forecast the effects of gene editing. To this end, we designed three levels of virtual and experimental mutation assays: *i*) motif insertion in vitro using tobacco leaf transient expression, *ii*) in silico motif insertion and single-nucleotide saturation mutagenesis, and *iii*) in vivo CRISPR-Cas9 editing in soybean (**Fig. 4)**. We first evaluated motif-scale edits by inserting cis-elements into the promoter of a highly expressed gene, *GmW82.01G042700*. Using *GenoRetriever*, we predicted the effect of randomly inserting a TATA-box at −143 bp and a YY1 motif at −220 bp and +18 bp relative to the TSS (**Fig. 4a)**. All three insertions were predicted to down-regulate TSS abundance (**Fig. 4a)**. In tobacco leaf transient assays (luciferase reporter), the observed expression changes closely matched *GenoRetriever*’s predictions, confirming that the model reliably anticipates motif-level editing outcomes (**Fig. 4a)**. Next, we assessed *GenoRetriever*’s resolution at the single-base level by simulating every possible nucleotide substitution and motif knockout within 500 bp upstream of the translation start site (ATG) (**Fig. 4b)**. For each variant, we computed the predicted relative change in the abundance of transcription initiation (**Fig. 4b)**. Tobacco transient assays validated that *GenoRetrieve*r correctly predicted the direction of expression change (up- or down-regulation) for single-nucleotide edits, demonstrating base-pair resolution in its forecasting capability (**Fig. 4b)**. Finally, we tested *GenoRetriever*’s utility for guiding real-world gene editing. We designed CRISPR-Cas9 guide RNAs targeting the 5′ UTR of a model gene and used the model to predict relative changes in TSS abundance for the edited alleles (**Fig. 4c, Supplementary Table X)**. Subsequent soybean transformations and STRIPE-seq profiling showed an 88.23% accuracy in predicting the direction of expression change, underscoring the model’s practical applicability despite the complexities of in vivo editing (**Fig. 4c)**. Together, these multi-scale validation experiments confirm that *GenoRetriever* can guide promoter engineering, from motif insertion to single-base edits and full in vivo gene editing, with high fidelity. To facilitate broader use, we have launched a free online service at xtlab.hzau.edu.cn/GenoRetriever. Users can upload promoter sequences to *i*) analyze motif effects, *ii*) predict expression changes in real time after in silico edits, and *iii*) perform saturated mutagenesis of the ±100 bp region around the TSS **(Extended Data Fig. 4**). We anticipate that this resource will accelerate intelligent design in crop improvement and functional genomics.

**Figure 4.**
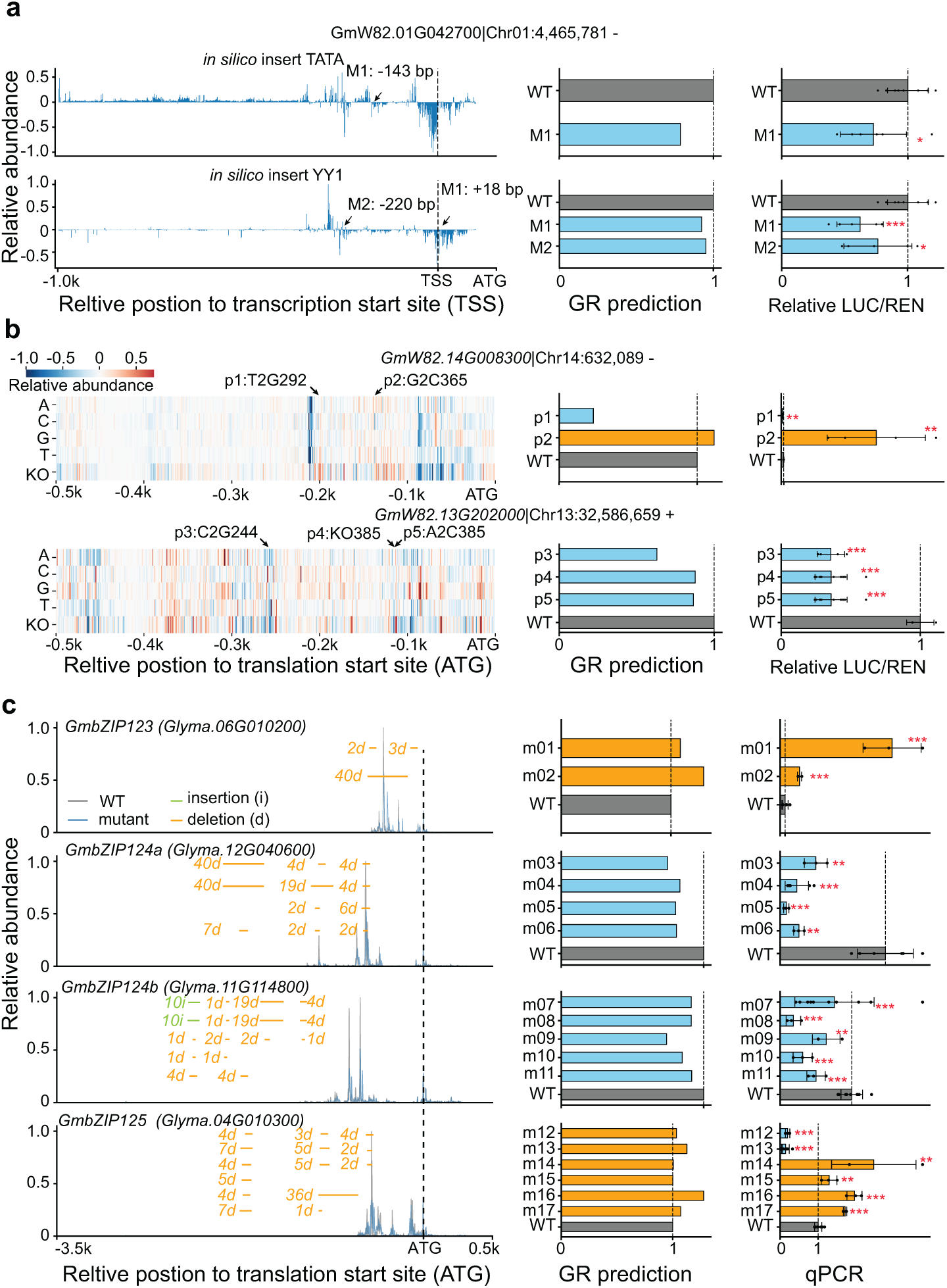
Validation of *GenoRetriever* predictions across multi-scale sequence edits. **a** Motif insertion effects for gene *GmW82.01G042700*. Left panel shows *GenoRetriever*’s *in silico* prediction of relative effect when a TATA-box or YY1 motif is inserted within 1-kb upstream of the translation start site (ATG). Arrows mark the insertion sites of each sample. Right panels compare predicted relative expression (middle panel) with tobacco leaf transient assay results (right panel). The y axis lists sample treatments and the x axis shows relative luciferase/renilla (LUC/REN) signal. **b** Single nucleotide saturation mutagenesis effects. Left panel is a heatmap of predicted changes for every possible nucleotide substitution within 500 bp upstream of the ATG. Rows represent mutation types and columns represent position relative to the ATG. Arrows mark the insertion sites of each sample. Right panels compare predicted relative expression (middle panel) with tobacco leaf transient assay results (right panel). The y axis lists sample treatments and the x axis shows relative LUC/REN signal. **c** 5′ UTR editing effects on transcription initiation. Left panel is a line plot of *GenoRetriever*’s predicted average relative TSS signal for each CRISPR-Cas9 edit at the 5′ UTR. Arrows mark each edit site. Right panels compare predicted relative expression (milddle panel) with the qPCR result of oybean editing assay (rigth panel). The y axis lists mutant constructs and the x axis shows relative expression levels. Student’s t test significance is indicated (\**p* < 0.05, \*\**p* < 0.01, \*\*\**p* < 0.001).

### Divergence of motif effects on transcription initiation between monocots and dicots

To broaden our understanding of transcription initiation across plant genomes and validate the robustness of *GenoRetriever*, we applied the same modified STRIPE-seq protocol and analysis pipeline to leaf tissue from six major crops: cotton (TM-1), rapeseed (ZS11), wheat (Svevo), tomato (LA1589), rice (ZS97), maize (B73) **(Supplementary Table 1).** We first used the soybean-trained consensus model of *GenoRetriever* to extract key sequence patterns from each species’ STRIPE-seq data. All patterns identified in these diverse crops fell within the set discovered in soybean, indicating that fundamental sequence determinants of transcription initiation are broadly conserved. We then evaluated three training strategies, direct application of soybean model weights, fine-tuning of soybean weights on each species’ data, and *de novo* training per species. Direct application achieved satisfactory performance (average Pearson correlation *r* > 0.65) in most crops, reflecting shared regulatory logic (**Fig. 5a)**. Fine-tuned models outperformed direct predictions, revealing subtle species-specific differences, and also surpassed *de novo* models, demonstrating the value of high-quality, multi-tissue soybean data for enhancing single-tissue predictions in other crops (**Fig. 5a, b)**. To investigate evolutionary divergence in motif regulation, we performed hierarchical clustering on the average relative effect of each motif in promoters across the eight species (**Fig. 5c, Supplementary Table 9)**. This analysis cleanly separated monocot and dicot species into distinct clusters, confirming that motif-to-TSS regulatory patterns differ subtly between these lineages (**Fig. 5c, Supplementary Table 9)**. We further quantified motif occurrence frequency and positional distribution relative to the TSS, finding a positive correlation (*r* = 0.70) between changes in motif effect and changes in motif frequency across monocots and dicots (**Fig. 5c, Supplementary Table 9 &10)**. Kernel density estimation with increased bandwidth (adjust = 5) revealed that some motifs, like the TATA-box, have narrow high-frequency intervals, whereas others, such as TCP20, display broader intervals (**Fig. 5d, Supplementary Table 9)**. Notably, shifts in the distance of motifs from the TSS were negatively correlated (*r* = −0.71) with changes in motif effect, suggesting that during evolution, key motifs often move closer to the TSS and increase their promoter frequency to enhance transcription initiation activity **(Supplementary Table 10)**.

**Figure 5.**
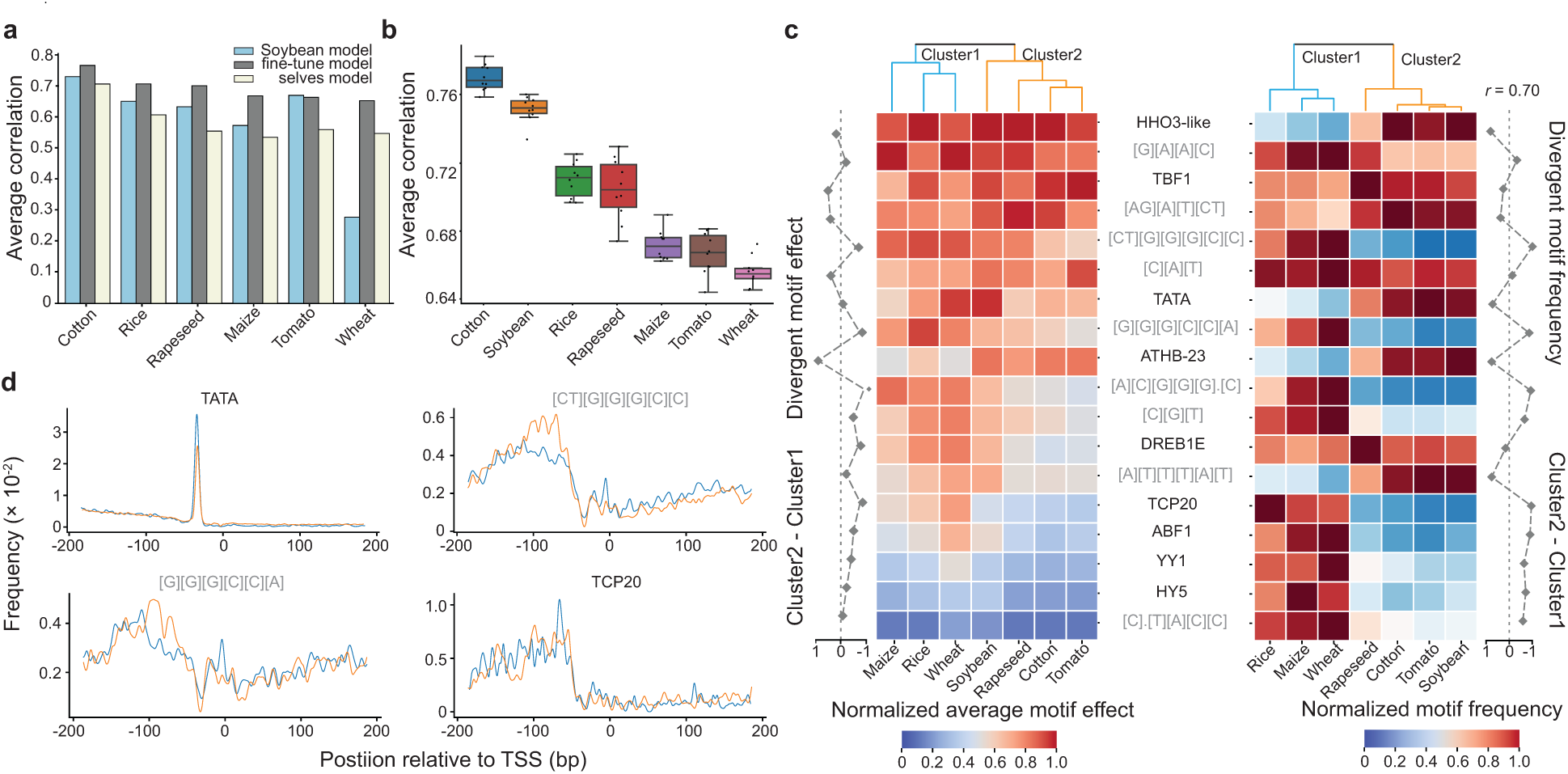
Evolutionary divergence of motif regulatory patterns on transcription initiation across plant species. **a** Comparison of three training strategies for predicting TSS signals in each plant, including direct application of the soybean model (blue), fine-tuning of the soybean model on each species’ data (gray), and *de novo* training on each species’ data (yellow). The y-axis shows the average Pearson correlation on the test set and the x-axis lists the plant species. **b** Prediction accuracy of fine-tuned *GenoRetriever* models across seven plant species. The y-axis shows the average Pearson correlation coefficient between predicted and observed TSS signals for each species. The x-axis lists the plant species. **c** Clustering of motif regulatory patterns across species. Left heatmap shows the average relative contribution of each motif to TSS signal for each crop, clustered by similarity. Right heatmap shows the average frequency of each motif in all promoters of each plant, clustered separately. The adjacent line plot depicts the difference in average motif effect between the two main clusters (monocots vs. dicots). **d** Kernel density estimates of motif positional distributions relative to the TSS for monocot and dicot species. The y-axis shows motif occurrence frequency and the x-axis shows position relative to the TSS (0 marks the TSS).

## Discussion

In this study we developed GenoRetriever, an interpretable deep learning model that predicts transcription initiation in plant genomes with high accuracy. We trained the model on modified STRIPE-seq data from eight soybean tissues (wild and cultivated) and from three monocot and three dicot species. A two-stage consensus network distilled 27 core sequence patterns, including both canonical motifs and initiator elements, and *GenoRetriever* learned each pattern’s contribution to transcript abundance and the precise location of transcription start sites. We validated these predictions through virtual motif insertions, *in silico* single-nucleotide saturation mutagenesis and CRISPR-Cas9 promoter edits, majority of which yielded expression changes in close agreement with our model forecasts. An *in silico* degradation experiment further showed that roughly 31.85% percent of natural promoter variation between wild and cultivated soybean drives shifts in dominant motifs during domestication. Finally, by building individual *GenoRetriever* models for each of the seven crop species, we confirmed that core sequence determinants remain conserved while monocots and dicots exhibit distinct motif-to-TSS regulatory patterns driven by differences in motif frequency and proximity to the TSS.

The consensus network architecture of *GenoRetriever* adapts ideas from the *Puffin* model^14^, originally developed for human transcription start modeling, but introduces key modifications for plant genomes. Instead of small convolution kernels of size three, we employ much larger kernels of 51 base pairs for initial motif sampling and 601 base pairs for deeper feature extraction, which capture longer and more complex plant cis-elements such as trinucleotide initiators. Pattern discovery and signal prediction are separated into two stages, with motif and supplementary effect networks first identifying robust, cross-tissue sequence patterns and then freezing those weights in the prediction network. To weight these features without adding positional noise, a Multi-head Efficient Channel Attention module assigns importance scores to feature channels rather than sequence positions. The downstream predictor uses a U-net style encoder-decoder with strided convolutions replacing pooling layers, linear interpolation instead of nearest neighbor upsampling and skip connections to combine local detail with global context across a 4-kb region. Training minimizes an L1 Pseudo Poisson Kullback-Leibler (KL) loss to balance count fidelity with sparsity, and a cosine annealing schedule improves generalization. We validated *GenoRetriever* on eight soybean tissues and seven other crop species through *in silico* ablation, tobacco leaf transient assays and in vivo CRISPR-Cas9 edits, preserving interpretability while extending predictive power to the evolutionary and structural complexity of plant transcription initiation.

Despite high resolution TSS profiling in *Arabidopsis*^20^, soybean^10^, cotton and maize^4^, our understanding of plant transcription initiation has remained limited to a few core cis-elements such as the TATA-box and initiator. Many plant promoters lack a TATA-box yet still initiate transcription efficiently, indicating additional uncharacterized motifs at work^10^. In the human^9^ genome, bidirectional promoters are common and linked to enhancer RNA production, but they are rare in both monocots and dicots^4, 14, 21^*. GenoRetriever* analysis distilled 27 motifs across seven plant species but found only one bidirectional element, DREB1E, providing a new explanation for the scarcity of bidirectional transcription start regions in plants. *GenoRetriever* interprets motif contributions to both transcript abundance and precise TSS positioning. Motif-driven changes in abundance shape expression patterns, while shifts in motif placement can create alternative transcription start sites that yield distinct protein isoforms in different tissues. We validated predictions for five well-characterized motifs using soybean hairy-root transfromation assays to minimize phenotypic shock. Although trans-acting factors introduce noise, the strong correlation between predictions and experimental results confirms that *GenoRetriever* captures the essential cis-regulatory code at single-base resolution. Comparative models for seven crops revealed that while core sequence patterns recur, their frequency and distance from the TSS differ by lineage, suggesting that evolutionary shifts in motif content and localization underlie lineage-specific regulatory strategies.

Prediction and interpretation of sequence determinants are fundamental to the precise design of promoters and other noncoding elements for controlled gene expression. To enable broad adoption, we launched an intuitive web server that integrates all eight species models with their STRIPE-seq datasets. Users can explore high resolution TSS profiles for any supported species, upload custom promoter sequences for *in silico* motif insertion or single-base mutagenesis and instantly view predicted effects on transcription initiation. Although we have validated editing predictions experimentally in soybean, the server currently supports only leaf tissue data for other crops. Researchers who generate STRIPE-seq data from additional tissues can retrain or fine-tune models using the same workflow described here, ensuring tissue-specific prediction accuracy. From an application perspective, precise promoter editing offers a stable alternative to transgene overexpression, which often loses efficacy after a few generations. By combining *GenoRetriever*’s accurate forecasts with advanced gene editing tools such as CRISPR base editing, prime editing or CRISPR activation and repression systems, researchers can fine-tune gene expression without creating knockout mutations. This integrated approach brings us closer to realizing the promise of precision breeding and functional genomics across multiple crop species.

## Materials and Methods

### Plant growth conditions and material collection

Wild soybean (*Glycine soja* PI468916) and cultivated soybean (*G. max* cv. Williams 82) were grown in a greenhouse under a 16 h light/8 h dark photoperiod at 25 ± 2 °C. Plants were maintained on a vermiculite:soil mix (3:1) and watered as needed. For nodulation assays, seedlings were inoculated with *Bradyrhizobium diazoefficiens* USDA 110 and grown for 20 days post-inoculation (dpi) alongside uninoculated controls, following the protocol of of Ren et al. (2019)^22^. Eight tissues were harvested at the development stages of trefoil, flowering, and seed development. Cotton (Gossypium hirsutum TM 1), rapeseed (Brassica napus ZS11), wheat (Triticum durum ‘Svevo’), tomato (Solanum lycopersicum LA1589), rice (Oryza sativa ZS97) and maize (Zea mays B73) were grown under the same greenhouse conditions. Leaf tissues were harvested at young developmental stages. For each species and tissue, samples were collected from at least eight individual plants, pooled, flash-frozen in liquid nitrogen, and stored at −80 °C until RNA isolation.

### RNA isolation and rRNA depletion

Total RNA was extracted from each tissue using TRIzol reagent (Invitrogen/Life Technologies, CA) following the manufacturer’s protocol. To eliminate genomic DNA contamination, RNA samples were treated with the TURBO DNA-free Kit (Invitrogen). For nodule tissue of PI468916, rRNA was removed in two steps: first with the NEBNext rRNA Depletion Kit (Bacteria) (New England Biolabs), then with the RiboMinus Plant Kit for RNA-Seq (Invitrogen). RNA from the remaining seven tissues underwent rRNA depletion using only the RiboMinus Plant Kit for RNA-Seq.

### Modified STRIPE-seq library construction and sequencing

To enrich for capped transcripts, total RNA was subjected to mRNA capture using VAHTS mRNA Capture Beads 2.0 (N403, Vazyme). The captured mRNA was then incubated with XRN-1 (R7040M, Beyotime) for one hour to digest non-capped RNAs. Template-switching reverse transcription (TSRT) (2964768, invitrogen) was then performed using a unique barcoded reverse transcription oligonucleotide (RTO) for each sample. A 5′-biotin modified, unique molecular identifier–containing template-switching oligonucleotide (TSO) with three 3′ riboguanosines was added to capture the 5′ end. VAHTS DNA Clean Beads (N411, Vazyme) cleanup is then performed to remove excess TSO and RTO. The resulting products were amplified by PCR using TruSeq adapter–compatible primers. Depending on the input RNA amount, 16 to 20 cycles of PCR were carried out with the forward library oligo (FLO) and reverse library oligo (RLO). Solid-phase reversible immobilization bead (N411, Vazyme) cleanup was used to size-select fragments between 250 and 750 base pairs. Final libraries (25-100 ng) were sequenced on an Illumina NovaSeq PE150 platform to produce 150-base-pair paired-end reads at Novogene Corporation Inc. (Tianjin, China). Primer sequences for the RTO, TSO and library PCR primers are listed in **(Supplementary Table 11)**.

### Processing of STRIPE-seq data

Raw sequencing read quality was assessed with FastQC version 0.11.9 (https://www.bioinformatics.babraham.ac.uk/projects/fastqc/). Only Read 1, which corresponds to the sense strand of RNA, was used for downstream analysis. Adapter and poly-G tails were removed with cutadapt version 2.5 (Martin 2021)^23^. Reads shorter than 50 nucleotides or lacking the TATAGGG motif were discarded. PCR duplicates were collapsed using the UMI information via the fastx_toolkit function fasx_collapser version 0.0.14 (http://hannonlab.cshl.edu/fastx_toolkit). The TATAGGG motif was then trimmed from the remaining reads with cutadapt. Cleaned reads were first aligned to rRNA sequences using HISAT2^24^ with default settings. Unmapped reads from this step were retained and then aligned to the reference genome sequences of each species^13^ with HISAT2 **(Supplementary Table 1)**. Only alignments with mapping quality greater than 30 were kept by filtering with SAMtools version 1.8^25^. For nodule samples, additional filtering removed any reads that mapped to the Bradyrhizobium diazoefficiens USDA 110 genome. Transcription start site regions (TSRs) were then identified, quantified, and annotated following the pipeline described in Wang et al. (2024)^10^, and all custom scripts are available on our GitHub repository (https://github.com/Wanjie-Feng/STRIPE_seq).

### Data preprocessing of *GenoRetriever*

For the single-tissue GenoRetriever model, transcription start sites (TSS) with counts per million (CPM) greater than one were selected as training examples. In the multi-tissue combined model, each TSS abundance was defined as the sum of CPM values across all tissues. To minimize false positives, only those TSS that exceeded a CPM of one in at least two tissues were retained for training.

Before input into the model, all CPM values were transformed using a log base two scale according to the formula

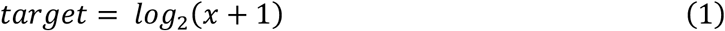

where x is the raw CPM.

After log transformation, the signals were smoothed using a one-dimensional convolution layer with a window size of five. This smoothing step reduces differences in apparent peak width between TSS and removes positional noise arising from experimental variation. For validation in other crop species, TSS were ranked by relative expression and the top 60% were used as training data. This selection excludes low-abundance sites that may represent experimental noise.

### Motif consensus network

The motif consensus network consists of two convolution layers. Considering the average motif length and common motif-mining window sizes, the first convolution layer uses a kernel size of 51 to sample sequence features (Supplemtnary Figure 2, Consensus network Stage 1). To preserve the raw features extracted by this layer, we apply only a Sigmoid activation function and omit batch normalization. The second convolution layer uses a kernel size of 601 to enhance feature extraction depth. After this layer, we apply batch normalization followed by a Softplus activation function to accelerate model convergence. The supplementary effect consensus network employs a two-branch, two-layer architecture (Supplemtnary Figure 2, Consensus network Stage 2). The first branch incorporates the distilled motif consensus patterns and their corresponding feature curves. During training, this branch’s convolution weights remain fixed; only the bias terms are updated. The second branch mirrors the motif consensus network structure with two convolution layers, but uses smaller kernels: size 3 in the feature extraction layer and size 15 in the feature encoding layer. All weights in this branch are trainable. We use Kullback-Leibler divergence^26^ as the primary loss function and include a PseudoPoissonKL regularization term with a scaling factor of 1 × 10^−3^ to balance trend extraction and abundance fidelity. To improve the visual clarity of the learned feature curves, we apply L2 regularization to the weights of the second convolution layer in each consensus network with a scale factor of 2 × 10^−3^. To prevent attention bias in downstream prediction, we apply L1 regularization to all trainable convolution weights: a scale factor of 4 × 10^−5^ for the first encoding layer and 5 × 10^−5^ for the second convolution layer.

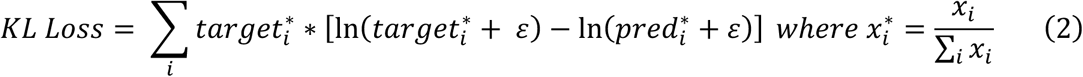

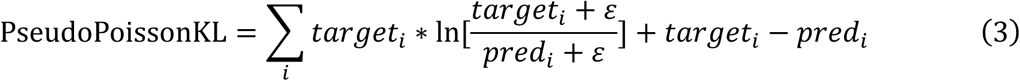

### Prediction network

The prediction network builds on the consensus networks to focus on accurate forecasting of transcription initiation signals. Its first layer contains two convolution modules whose weights are initialized from the motif consensus network and the supplementary effect consensus network, respectively, via knowledge distillation. During model training, the weight parameters of these two convolution modules remain fixed; only their bias terms are updated to allow fine adjustments. Following these fixed convolution layers, the network employs a U-net-style encoder-decoder architecture in which pooling layers are replaced by strided convolutions and nearest-neighbor upsampling is replaced by linear interpolation. Skip connections link corresponding encoder and decoder layers, facilitating the fusion of features at different scales. Within each encoder block, a feature extraction module first applies a multi-head efficient channel attention mechanism to reweight channel features, and then uses a bottleneck-style feedforward structure. The bottleneck feedforward structure expands and compresses feature dimensions through linear layers and integrates a convolutional hidden layer. Outputs from the attention mechanism and the feedforward structure are merged via residual connections to ensure stable gradient flow and capture both local and global sequence features. Two such U-net modules are cascaded, and a SoftPlus activation function at the end produces the predicted transcription initiation signal. During training, the model uses an L1-PseudoPoissonKL loss function to balance sparsity and count distribution realism. A cosine annealing learning rate schedule accelerates model fitting, prevents convergence to local minima, and enhances generalization performance.

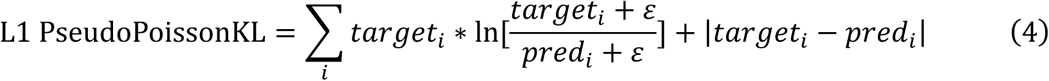

### Multi-head efficient channel attention (MECA)

The prediction network relies on motifs and supplementary sequence patterns distilled by the consensus networks, so its performance critically depends on accurately weighting these distinct feature channels. Traditional multi-head attention mechanisms emphasize positional relationships and are well suited to natural language, where tokens carry semantic roles. However, DNA sequences lack such clear “word” boundaries, and positional attention can introduce noise rather than clarity. To address this, we designed a channel-focused attention mechanism called Multi-head Efficient Channel Attention (MECA). MECA operates entirely along the channel dimension, ignoring sequence position to avoid misassigned attention. First, an adaptive average-pooling layer condenses each channel’s feature map into a single descriptor, capturing its global importance with minimal added parameters. The pooled descriptors are then split into multiple heads, each representing a distinct group of channels. Within each head, a lightweight linear projection network computes attention scores for its channels, enabling the model to learn diverse patterns of channel importance. Finally, the original feature maps are rescaled by these per-channel attention weights, enhancing the representation of critical sequence patterns while suppressing irrelevant noise. This approach delivers efficient and accurate channel weighting, boosting overall transcription initiation prediction performance.

### Artificial knowledge distillation

To identify robust sequence patterns across tissues and varieties, we applied an artificial knowledge distillation procedure based on convolutional kernel correlations. In the motif extraction stage, fifteen independent motif consensus models were trained separately on STRIPE-seq data from eight tissues of both Williams 82 and PI468916. For each tissue and variety, we computed pairwise Pearson correlations among all convolutional kernels in the feature-extraction layer. These correlations formed the edges of a connectivity graph, in which we retained only edges with correlation coefficients above 0.9. We then identified connected components whose size exceeded 30 percent of the number of trained models. Within each such component, the kernel (node) exhibiting the highest degree of connectivity was selected as its representative sequence pattern. After processing all tissues independently, we merged the resulting sequence patterns from both varieties and applied the same connectivity-based clustering procedure to the combined set. Following visualization of these merged patterns, we manually filtered out redundant or overly similar motifs. This workflow yielded a final, non-redundant collection of 27 distinct sequence features for downstream analysis.

### Ablation study

Ablation experiments were performed to dissect the functional contributions of individual sequence patterns encoded in *GenoRetriever*’s convolutional layers. In this model, each convolutional kernel in the encoding network represents a distinct sequence motif or initiator element and uses a non-negative Sigmoid activation. By setting the post-activation output of a given kernel to zero, we effectively silence that sequence pattern. The effect of this perturbation is quantified by comparing the predicted transcription initiation signal before and after silencing. We carried out three scales of ablation analysis: *i*) We simultaneously silenced all kernels corresponding to either motif patterns or Inr elements. By measuring the overall change in predicted TSS signal, we assessed the combined influence of each pattern class on transcription initiation. *ii*) Each encoding kernel was silenced one at a time. For each kernel, we calculated the change in predicted TSS abundance and its positional effect across a defined window around the TSS. This yielded individual abundance-effect and position-effect profiles for every motif and Inr pattern. *iii*) We simulated point mutations at each nucleotide position in the input sequence to identify critical variant sites. For every possible single-base substitution, we compared the change in predicted TSS signal. Variants causing the largest prediction shifts were flagged as key drivers of motif reconfiguration and transcriptional initiation changes, providing a robust approach for evolutionary analysis and regulatory target discovery.

### Motif effect scoring

To quantify the impact of each motif on transcription initiation, we computed two complementary scores based on the ablation experiment outputs. For each transcription start site (TSS), we obtained two signal tensors: one predicted before motif silencing and one predicted after silencing. We extracted a window of one thousand base pairs centered on the dominant position of the TSS (five hundred base pairs upstream and downstream). By subtracting the post-silencing tensor from the pre-silencing tensor at each position, we generated an effect curve whose values represent the magnitude of abundance change induced by the motif at single-base resolution. To capture both positional shifts and overall abundance changes in a single metric, we converted the same pair of 1-kb signal windows into the frequency domain using a discrete Fourier transform. In frequency space, we computed the pointwise squared difference between the two spectra and summed across all frequency components. The resulting scalar score reflects the combined positional and abundance effects of motif removal on transcription initiation.

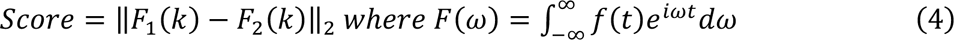

where *f(t)* is the time-domain signal (the effect curve), *F(ω)* is its frequency-domain representation, *ω* is the angular frequency, and iii is the imaginary unit.

### *In silico* degradation experiment and critical variation identification

To investigate how sequence variants affect transcription initiation signals between soybean varieties, we performed an in silico degradation experiment. For each promoter, we collected all variants including single-nucleotide polymorphisms and small indels which located within 2-kb upstream and 500 bp downstream of the transcription start site. We then ordered these variants in reverse evolutionary sequence and, one by one, reverted each derived allele back to the ancestral allele in silico. After each reversion, we recalculated the contribution score of every motif to the predicted TSS signal, using the metric defined in formula (4). A variant was designated as a critical variation if its reversion caused the promoter’s dominant motif (the motif with the highest contribution score) to switch from its original major type to a different motif type. This procedure yielded a set of critical variations that drive shifts in motif hierarchy and thus in transcription initiation regulation.

### T2T genome assembly and annotation

High-molecular-weight DNA was extracted using a CTAB-based protocol. For long-read sequencing, PacBio HiFi libraries were prepared and run on the Sequel IIe platform, while ultra-long read libraries for Oxford Nanopore Technologies were generated with the Ultra-Long DNA Sequencing kit and sequenced on PromethION. Hi-C libraries were constructed following Belton et al. (2012)^27^ and sequenced on a BGI DNBSEQ-T7 system. Short-read libraries were made with the NEBNext Ultra II DNA Library Prep Kit and sequenced on an Illumina platform. The telomere-to-telomere assembly was produced by integrating PacBio HiFi and ONT reads using hifiasm (v0.19.8-r603)^28^ and NextDenovo (v2.5.2)^29^. Contigs were screened against the NT database to remove plastid and ribosomal DNA sequences. Chromosome-level scaffolding used Hi-C data processed with Juicer (v1.6)^30^, refined through the 3D-DNA pipeline (v180922)^31^, and manually corrected in Juicebox (v1.9.8) (https://github.com/aidenlab/Juicebox). Assembly accuracy was confirmed by realigning HiFi and ONT reads and assessing completeness with BUSCO (v4.1.4)^32^ and Merqury (v1.3)^33^. Transposable elements were annotated using EDTA (v2.0.0)^34^, which integrates LTR_FINDER, TIR-Learner, and MITE-Hunter. Gene models were predicted through a combination of transcript evidence, homology, and ab initio methods. RNA-seq data from six tissues were aligned to the assembly with HISAT2 (v2.2.1)^24^ and assembled into transcripts using StringTie (v2.2.1)^35^. These models were refined by PASA pipeline (v2.5.3)^36^ and Braker3 (3.0.6)^37^, yielding a high-confidence gene annotation.

### Site-directed mutagenesis

Point mutations were introduced into recombinant plasmids by overlap extension PCR. Primers were designed with 15-20 bp overlaps flanking the mutation site. Each 10 µL PCR contained 50 ng template DNA, 0.5 µM of each primer and 2 × Phanta Flash Master Mix. Cycling conditions were 95 °C for 2 min; 30 cycles of 95 °C for 15 s, annealing from 52 °C (increasing by 0.4 °C per cycle) for 30 s, and 72 °C for 2 min; followed by a final extension at 72 °C for 5 min. PCR products were treated with DpnI at 37 °C for 1 h to remove template plasmid and then transformed into *E. coli* DH5α. Successful mutants were confirmed by Sanger sequencing. Primer sequences are listed in **(Supplementary Table 11).**

### Promoter Cloning and Dual Luciferase Reporter Assay

For dual luciferase assays, the regions upstream of the transcription start sites of *Glyma.01G042700* (970bp), *Glyma.13G202000* (665bp) and *Glyma.14G008300* (992bp) were amplified from Williams82 genomic DNA. Each fragment was cloned into the pGreenII 0800 LUC vector upstream of the firefly luciferase gene using KpnI (R3142, NEB) and BamHI (R3136M, NEB) restriction sites. Point mutations were introduced by overlap extension PCR. Primers were designed with 15 to 20 bp overlaps flanking each mutation site. Each 10 µL PCR contained 50 ng of recombinant plasmid DNA, 0.5 µM of each primer and 2 × Phanta Flash Master Mix (P510, Vazyme). Cycling conditions were 95 °C for 2 min; 30 cycles of 95 °C for 15 s, annealing from 52 °C increasing by 0.4 °C per cycle for 30 s, and 72 °C for 2 min; final extension at 72 °C for 5 min. PCR products were treated with 1 µL DpnI (R0176V, NEB) at 37 °C for 1h to digest parental DNA, then transformed into Escherichia coliDH5α (CC96102, TOLOBIO) competent cells. Mutations were confirmed by Sanger sequencing. Wild-type and mutant constructs were transformed into *Agrobacterium tumefaciens* strain GV3101. Agrobacterium cultures were grown to OD₆₀₀ = 0.6, harvested by centrifugation, and resuspended in infiltration buffer (10 mM MES pH 5.6, 10 mM MgCl₂, 100 µM acetosyringone). After incubating at room temperature for 2-3 hours, suspensions were infiltrated into the abaxial surface of *Nicotiana benthamiana* leaves. Plants were incubated for 48-72 hours under standard growth conditions. Firefly and Renilla luciferase activities were measured using a Tecan Spark microplate reader. Primer sequences are listed in **(Supplementary Table 11)**.

### Kowck down plasmid Construction and soybean hairy-root transformation

For RNA interference of the GmYY1 (Glyma.08G126300 and Glyma.05G167900), GmABF1 (Glyma.02G131700 and Glyma.07G213100), GmDREB (Glyma.09G147200, Glyma.10G239400, Glyma.16G199000 and Glyma.20G155100), GmHY5 (Glyma.18G117100 and Glyma.08G302500) and GmTCP (Glyma.16G053900 and Glyma.19G095300) gene families, conserved regions of 200 to 300bp were amplified from Williams82 genomic DNA and cloned into the pZHY930 vector using KpnI (R3142, NEB) and SmaI (R0141, NEB) restriction sites. Recombinant plasmids were validated by Sanger sequencing to confirm insert integrity and orientation. Final constructs were transformed into *Agrobacterium rhizogenesstrain* K599 for soybean hairy-root transformation. Transgenic roots were identified under a handheld lamp (3415RG, LUYOR) and knockdown efficiency was validated by RT-PCR with gene-specific primers. K599 carrying the desired constructs was used to generate soybean hairy roots following the protocol of Wang et al. (2014). Composite seedlings were transferred to pots (13 × 10 × 8.5 cm) filled with vermiculite and grown under a 16 h light / 8 h dark cycle at 25 °C and 50% relative humidity. Ten days after transfer, transformed roots were identified using a handheld lamp (3415RG, LUYOR). Transgenic roots were excised, immediately frozen in liquid nitrogen and stored at −80 °C until further analysis.

### CRISPR/Cas9 knockout vector construction

The CRISPR/Cas9 multiplex gene knockout vector pGES704 was employed to target the upstream open reading frames (uORFs) in the promoter regions of four soybean genes: *GmbZIP123* (*Glyma.06G010200*), *GmbZIP124a* (*Glyma.12G040600*), *GmbZIP124b* (*Glyma.11G114800*), and *GmbZIP125* (*Glyma.04G010300*). The pGES704 vector was derived from the base plasmid pGES401^38^, in which the selection marker gene *bar* was replaced with *CP4 EPSPS*. Six single-guide RNAs (sgRNAs) were designed in tandem to target these four genes (specific sequences and detailed information provided in **Supplementary Table 11**). Notably, some sgRNAs were engineered to simultaneously target 2-4 genes. The sgRNA-containing fragments were assembled with the vector using the NEBridge^®^ Golden Gate Assembly Kit (BsaI-HF^®^ v2), and successful plasmid construction was verified by Sanger sequencing.

### Transgenic plant generation and genotyping analysis

The validated knockout vectors were transformed into Agrobacterium tumefaciens strain GV3101 for stable soybean transformation via the cotyledonarynode transformation method as described^39^. Genomic DNA was extracted from T_2_ generation plants using the cetyltrimethylammonium bromide (CTAB) method, followed by PCR amplification with Taq Plus Master Mix II (P213, Vazyme). Target-specific primers for genotyping are listed in **Supplementary Table 11**. Mutation patterns of T_2_ generation plants were identified through Sanger sequencing.

### Quantitative RT-PCR analysis

Total RNA was isolated from leaves of T_2_ plants at the same developmental stage using the FastPure Universal Plant Total RNA Isolation Kit (RC411-01, Vazyme). Reverse transcription was performed with HiScript III RT SuperMix for qPCR (R323, Vazyme). Quantitative PCR was conducted using ChamQ Universal SYBR qPCR Master Mix (Q711, Vazyme) on a real-time PCR system (qTOWER3G, analytikjena, Germany). Three biological replicates, each with three technical replicates, were conducted for each sample. Gene expression levels were calculated using the 2^−ΔΔCt^ method^40^, with qPCR primers detailed in **Supplementary Table 11**.

## Data Availability

The data generated and reported in this study have been deposited in the GSA database (https://ngdc.cncb.ac.cn/gsa/) under accession number **PRJCA038998**. The genome assembly and related scripts used for data analysis are available upon request or can be accessed on GitHub at https://github.com/Wanjie-Feng/GenoRetriever. All TSS datasets generated in this study have been deposited on GitHub and are also available for download via the *GenoRetriever* web server.

## Acknowledgements

This work was primarily supported by the National Key Research and Development Program of China (2022YFD1201502, 2022YFD201400), the Natural Science Foundation of China (32272064, 32330078). We also thank Dr Xin Wang, Dr Handong Su, Dr Ting Zhao and Dr Hu Zhao for providing valuable tomato, wheat, rice, rapeseed and cotton seeds.

## Author Contributions

XW, PG conceived and designed the study. PG performed the primary data analysis and developed the GenoRetriever architecture and models. LL, LQ, XL, JY, ZL, JL, YG and XL carried out the experiments. WF and SH assembled the PI468916 T2T genome. YM, PG and HZ developed the *GenoRetriever* web server., XW and PG drafted the manuscript. All authors reviewed and approved the final version of the paper.

---

**Extended Data Fig. 1.**
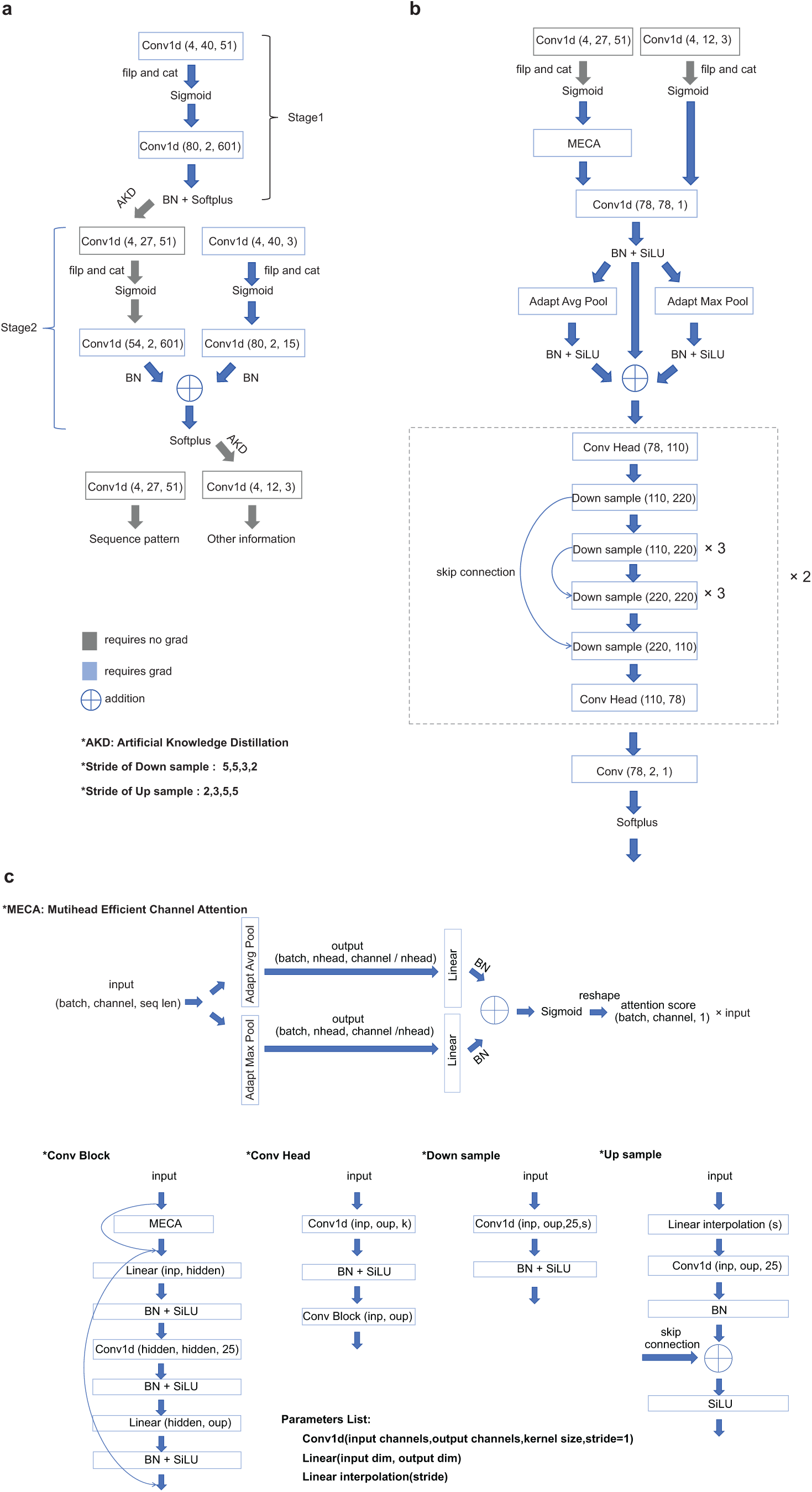
Architecture of the *GenoRetriever* deep learning model. **a** The consensus network extracts key information from genomic sequences. In stage 1 it identifies core sequence motifs, and in stage 2 it captures additional supplementary effects beyond those motifs. **b** The prediction network then uses these distilled features to accurately forecast transcription initiation signals during mechanistic analysis. **c** The architecture details of each module, including the specific layers, kernel sizes and attention mechanisms employed in each network component.

**Extended Data Fig. 2.**
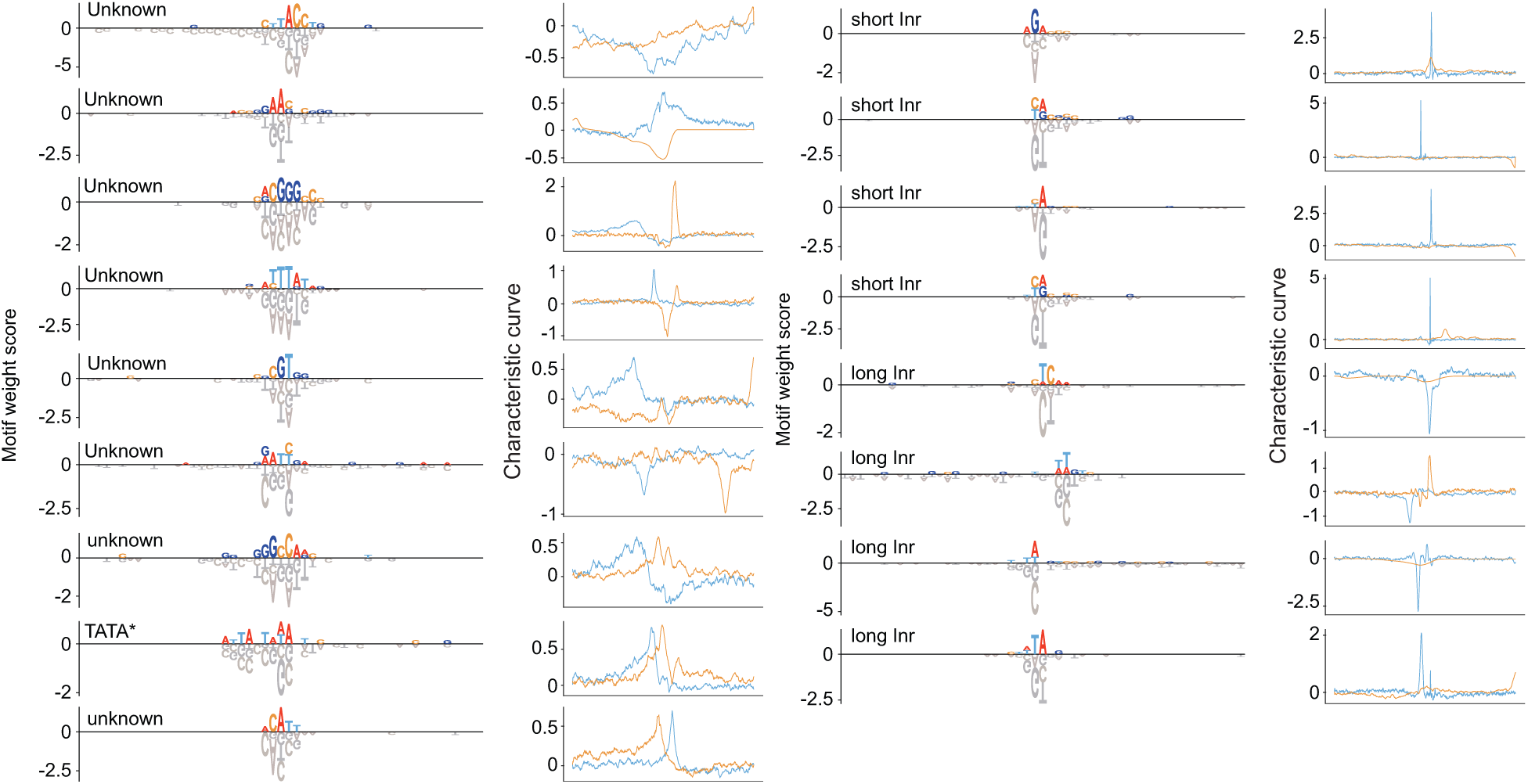
Characteristic response curves for nine novel motifs and eight Inr sequence patterns. Out of the 27 patterns identified by the motif consensus network, this figure displays the nine previously uncharacterized motifs (left panel) and eight Inr (right panel). Each panel shows the pattern logo and its average response curve over a 1 kb window centered on the pattern’s peak position.

**Extended Data Fig. 3.**
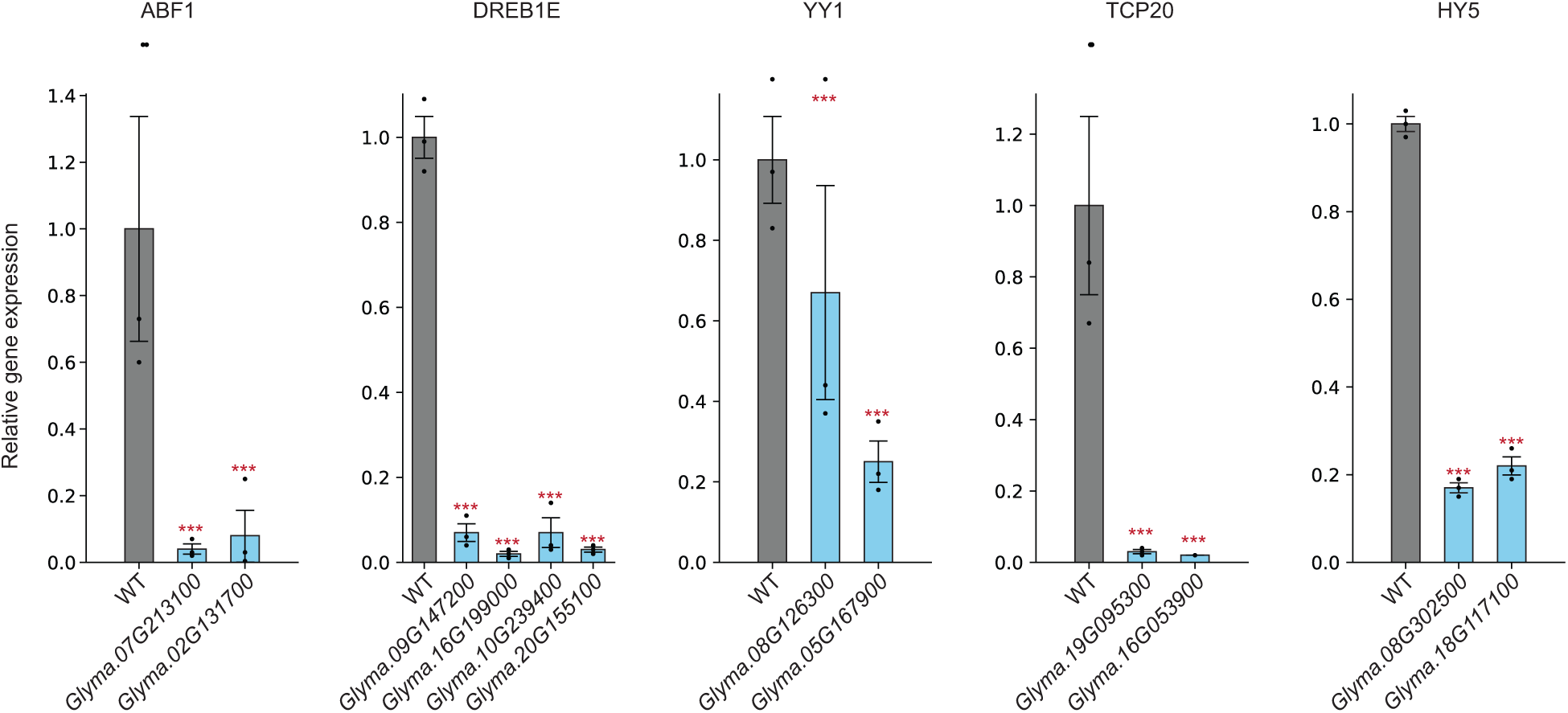
qPCR validation of gene knockdown efficiency in soybean hairy roots. Relative expression levels of target genes in RNAi lines compared to control roots. Data represent mean ± SD from three biological replicates. Statistical significance was determined by Student’s t-test (***p < 0.001).

**Extended Data Fig. 4.**
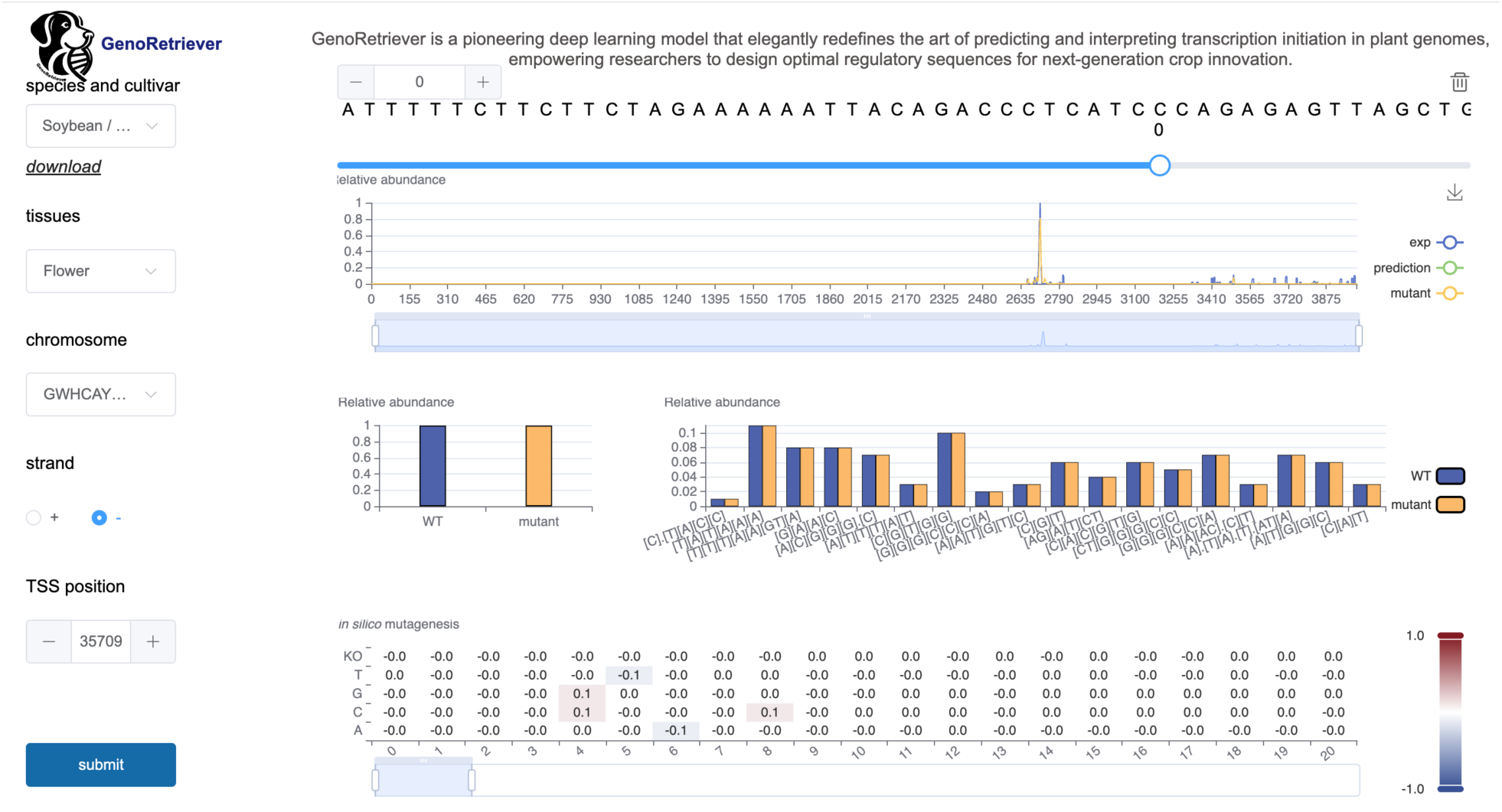
User interface of the GenoRetriever web server and module descriptions. The GenoRetriever web server provides an intuitive platform for exploring transcription start site profiles and predicting sequence edits.

## Supplemental information

**Supplementary Table 1. Sequencing metrics and TSS detection summary for all STRIPE-seq libraries.**

**Supplementary Table 2. Quantitative effect scores of each motif on transcription initiation across all Williams 82 promoters.** The motif index indicates the ordinal position of each motif within this table.

**Supplementary Table 3. Summary of sequencing data used for PI468916 genome assembly.**

**Supplementary Table 4. Genome assembly and annotation statistics for the PI468916 wild soybean accession.**

**Supplementary Table 5. Summary of transcription start sites (TSS) detected in multiple tissues of PI468916.** TSR indicates each defined transcription start region.

**Supplementary Table 6. Quantitative effect scores of each motif on transcription initiation across all PI468916 promoters.** The motif index indicates the ordinal position of each motif within this table.

**Supplementary Table 7. Paired promoter relationships between Williams 82 and PI468916, including syntenic coordinates.** TSR indicates each defined transcription start region.

**Supplementary Table 8. Key sequence variants driving motif shifts between Williams 82 and PI468916.** The motif index indicates the ordinal position of each motif within this table.

**Supplementary Table 9. Motif effect sizes, occurrence frequencies, and positional distributions across seven crop species.**

**Supplementary Table 10. Comparative changes in motif regulatory effects between monocot and dicot species, including effect size, positional distribution, and occurrence frequency.**

**Supplementary Table 11. List of gRNA sequences, primers, and adapter oligonucleotides used in this study.**

